# Unveiling the temporal impact: Exploring dynamic changes in the paediatric solid tumour immune microenvironment through time

**DOI:** 10.1101/2025.10.07.680325

**Authors:** Virgile Raufaste-cazavieille, Stéphanie Bianco, Lara Hermann, Jeyani George Clement, Emeric Texeraud, Sylvie Langlois, Thomas Sontag, Pascal Tremblay-Dauphinais, Alex Richard-St-Hilaire, Charles Joly Beauparlant, Damien Faury, Stéphanie Vairy, Nada Jabado, Sonia Cellot, Vincent-Philippe Lavallee, Thai Hoa Tran, Daniel Sinnett, Arnaud Droit, Raoul Santiago

## Abstract

The composition of the tumour immune microenvironment (TIME) influences tumour evolution and responsiveness to immunotherapy. While longitudinal changes in TIME have been well-characterized in adult cancers, its dynamics in childhood cancers remain poorly documented, limiting our ability to predict treatment responses and tailor immunotherapeutic strategies. This study aimed to evaluate the plasticity of TIME in paediatric solid tumours, investigate its longitudinal evolution, and identify time-dependent immune alterations. Transcriptomic data from longitudinal samples of 27 paediatric patients (<21 years old) with relapsed or refractory solid tumours were analysed, encompassing 70 timepoints: 16 diagnoses and 54 successive relapses. TIME plasticity was assessed using gene expression clustering and immune cell infiltration enumeration. Patient-adjusted longitudinal analyses were performed using generalised linear mixed models (glmmSeq), adjusted for age and sex. Temporal associations of immune changes were further explored using dynamic regression models. Thirteen patients exhibited significant changes in their TIME profile, indicating high TIME plasticity. Over time, the TIME shifted toward a tolerogenic and immunosuppressive state, characterised by decreased activity in immune pathways (e.g., T cell receptor signalling) and enrichment of tolerogenic (e.g., macrophage differentiation) and oncogenic pathways (e.g., IL6-JAK-STAT3). The core enrichment of upregulated pathways contained key immunosuppressive factors: immune checkpoints (CTLA-4), tumour-associated macrophage activators (CSF1/CSF1R), T-regulatory cell activators (TGFB1), and immunosuppressive genes (IL10RA). This study provides evidence that the TIME in paediatric solid tumours is plastic and remodels towards immune depletion and tolerogenicity. This evolution may underlie treatment resistance and disease progression, underscoring the need for TIME-informed therapeutic approaches in paediatric oncology.

**Significance Statement:** This article demonstrates the plasticity of the tumour immune micro-environment (TIME) of paediatric solid tumours throughout disease evolution. Longitudinal transcriptomic analyses of 70 tumour samples from 27 patients showed a progressive remodelling towards tolerogenicity and immune depletion. Key immunosuppressive factors, including immune checkpoints and tumour-associated macrophages, were identified as potential contributors to immune escape. These findings support the relevance of longitudinal immune monitoring in paediatric oncology and may inform future strategies for immunotherapeutic interventions.

## Introduction

Cancer remains the leading cause of disease-related mortality among children in high-income countries (1). While immune checkpoint blockade (ICB) therapies have demonstrated significant efficacy in adult cancers (2), their success in paediatric solid tumours has been limited, with response rates reported in only ∼3% of cases (3). Understanding the mechanisms underlying resistance to ICBs is essential for improving patient stratification and optimizing immunotherapeutic strategies in paediatric oncology (4).

The effectiveness of ICB is closely linked to the tumour immune microenvironment (TIME). In adult cancers, several biomarkers—including high PD-L1 expression, elevated tumour mutational burden (TMB), and a T cell-inflamed phenotype—are associated with improved responses to ICB (5,6). Conversely, the presence of immunosuppressive elements such as tumour-associated macrophages (TAMs), regulatory T cells (Tregs), and low TMB are linked to resistance. These predictive markers are less reliable in paediatric cancers. Most paediatric tumours exhibit low TMB and minimal PD-L1 expression, and the correlation between PD-L1 transcript and protein levels remains poorly defined. Notably, stratification based on PD-L1 expression alone has proven insufficient to identify likely responders to ICB in paediatric cohorts (7). Novel immune-based classification of paediatric tumours’ immune environment could distinguish distinct immune phenotypes: hot, altered, and cold (8). Despite the rarity of children treated with immunotherapy for solid tumours, a recent work identified markers associated with response to ICB specific to paediatric cancers. High CD8+ T cell infiltrate, high PD-L1 expression, and TCR diversity were associated with improved progression-free survival (9). This work reinvigorated the capacity to identify childhood cancers that can benefit from immunotherapy when tailored to their molecular specificity. However, if biomarkers can guide targeted immunotherapy, a major limitation of current biomarker-driven approaches is the assumption that the TIME is a static entity and remains stable during disease evolution.

Both intrinsic (tumour clonal evolution) and extrinsic factors (treatment) can influence the TIME composition. This immune plasticity has been well documented in adult cancers. For example, in pancreatic cancer, when comparing different invasive stages, the T cell populations decline during tumour progression, while immunosuppressive populations increase, revealing evolution of TIME due to intrinsic factors (10). After cancer treatment, chemotherapy induced T cell exclusion in breast cancers (11), while ICB combination (anti-PD-1 and anti-CTLA-4) enhanced T cell clonality in melanoma, demonstrating treatment-related TIME evolution in adult malignancies (12).

In paediatric cancer, the plasticity of TIME hasn’t yet been studied, and its temporal dynamics remain an uncharted area. Addressing this gap is instrumental for understanding the drivers of ICB resistance and determining the best strategies and optimal timing for integrating ICB in the therapeutic options for paediatric cancers. In this study, we hypothesized that the TIME in paediatric solid tumours is plastic and tends to evolve toward a more tolerogenic state over time. To test this, we conducted the first longitudinal analysis of the TIME in paediatric cancers, leveraging a cohort of 27 patients and 70 tumour samples collected across multiple timepoints during disease progression. Our goal was to determine whether the TIME remains stable or undergoes significant changes, and to characterize the directionality of these changes—either toward immune activation or suppression. This work aims to provide foundational insights into TIME evolution in paediatric cancers to help guide clinicians in selecting patients who may benefit from immunotherapy.

## Results

### Population description

From the SIGNATURE database, the oncogenomic initiative of the province of Quebec, we identified 27 patients with disease recurrence and at least two timepoints collected. This longitudinal cohort consisted of 15 males and 12 females, with 2 patients diagnosed with central nervous system (CNS) and 25 with extracranial solid tumours (Supplementary Table S1). The main tumour types were: 6 osteosarcomas, 4 neuroblastomas, 3 rhabdomyosarcomas, 5 other soft tissue sarcomas, 2 high-grade gliomas, 2 carcinomas, and 5 other unique malignancies. Sample collection timepoints included a total of 70 samples: 16 at initial diagnosis, 22 at first relapse, 18 at second relapse, 14 at third or subsequent relapses. At the time of biopsy, 7 patients were < 2 years of age, 15 from 2 to < 6 years, 17 from 6 to < 12 years, and 31 from 12 to < 21 years.

### The tumour immune microenvironment is plastic in solid paediatric cancer

To identify if paediatric tumours were plastic over time, we introduced the concept of “immune distance” to assess temporal changes in the tumour immune environment. The goal was to distinguish patients with significant immune changes over time. Immune distance was determined by two metrics: immune phenotype (classified as hot or cold) and immune gene expression clustering.

The immune phenotyping, trained on 340 paediatric solid tumours from the SIGNATURE database, revealed that some patients with multiple disease occurrences switched from hot to cold, or vice versa, while others remained stable through recurrences (Figure 1A). For instance, patient 19 exhibited a consistent cold phenotype at all three timepoints, indicating stable immune gene expression. In contrast, patient 25 demonstrated substantial divergence between timepoints, with a cold phenotype at first and second relapses and a hot phenotype at the third relapse, suggesting dynamic changes of the immune environment over time. In total, six patients underwent a switch of immune phenotype during disease evolution (Table 1).

**Table 1.** Classification of the patients for the two metrics defining the immune distance. The table summarises the metrics for the different timepoints of each patient. Each timepoint is characterised with its immune phenotype (cold or hot) and immune cluster (1 to 5). The immune distance recapitulates the changes of the two metrics to characterise the immune plasticity/stability for each patient (distant or close) Abbreviations: D: diagnosis, ID: identification, R: relapse, tho: thorax, met: metastasis, nod: nodule

| Patient | Immune phenotype | Centroid | Pathology | Group |
| --- | --- | --- | --- | --- |
| Patient 1 | Cold | R1: 2, R2: 1 | Hepatocarcinoma | Distant |
| Patient 2 | Cold | R1/R6-tho: 3, R2/R5-mass/R5-met: 5, R6-nod:2 | Osteosarcoma | Distant |
| Patient 6 | Cold | R1/R3: 2, R2:3 | Soft tissue sarcoma | Distant |
| Patient 10 | R8: Hot, R1/R3/R6: Cold | D/R3/R6/R8: 3, R1: 5 | Soft tissue sarcoma | Distant |
| Patient 11 | Cold | R1: 2, R2: 3 | Osteosarcoma | Distant |
| Patient 13 | Cold | D/R2: 3, R1 : 5 | Osteosarcoma | Distant |
| Patient 14 | Cold | R1: 2, R2: 3 | Osteosarcoma | Distant |
| Patient 18 | D : Hot, R1 : Cold | D: 3, R1: 2 | Neuroblastoma | Distant |
| Patient 20 | Cold | D: 4, R: 2 | Rhabdoid tumour | Distant |
| Patient 21 | R3 : Hot, R2 : Cold | All timepoint in 5 | Osteosarcoma | Distant |
| Patient 22 | D : Hot, R2 : Cold | All timepoint in 2 | Neuroblastoma | Distant |
| Patient 23 | R1: Hot, R2 : Cold | R1: 3, R2: 2 | Osteosarcoma | Distant |
| Patient 25 | R3: Hot, R1/R2: Cold | R1/R2: 2, R3:3 | Neuroblastoma | Distant |
| Patient 3 | Cold | All timepoints in 2 | Soft tissue sarcoma | Close |
| Patient 4 | Cold | All timepoints in 2 | Carcinoma | Close |
| Patient 5 | Cold | All timepoints in 2 | Pleuropulm blastoma | Close |
| Patient 7 | Cold | All timepoints in 5 | Rhabdomyosarcoma | Close |
| Patient 8 | Cold | All timepoints in 3 | Soft tissue sarcoma | Close |
| Patient 9 | Cold | All timepoints in 3 | Rhabdomyosarcoma | Close |
| Patient 12 | Cold | All timepoints in 2 | Carcinoma | Close |
| Patient 15 | Cold | All timepoints in 2 | Germ cell tumour | Close |
| Patient 16 | Cold | All timepoints in 2 | Wilms tumour | Close |
| Patient 17 | Cold | All timepoints in 2 | Rhabdomyosarcoma | Close |
| Patient 19 | Cold | All timepoints in 2 | Soft tissue sarcoma | Close |
| Patient 24 | Cold | All timepoints in 2 | Neuroblastoma | Close |
| Patient 26 | Cold | All timepoints in 2 | High-grade glioma | Close |
| Patient 27 | Cold | All timepoints in 2 | High-grade glioma | Close |

**Figure 1.**
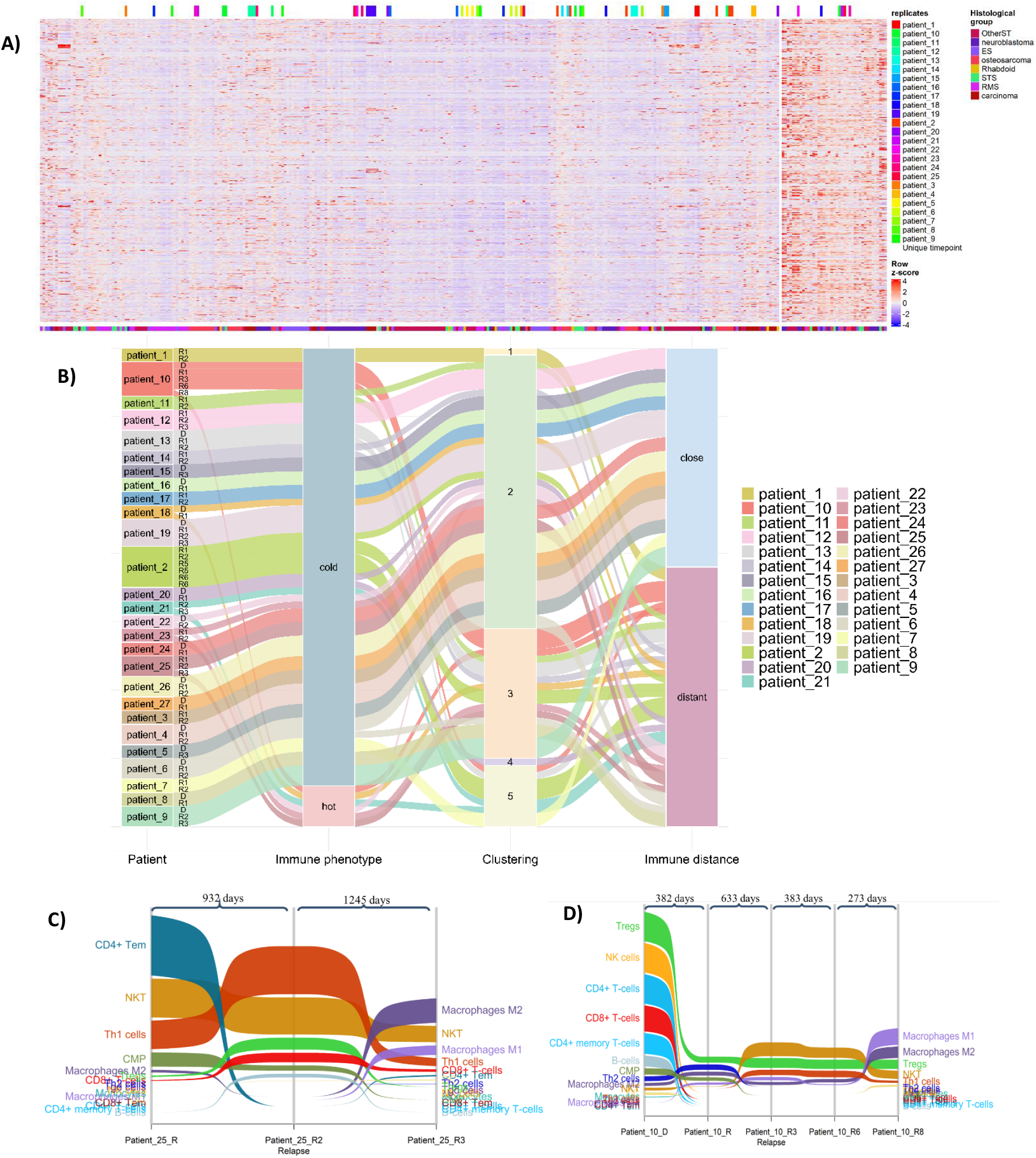
Representation of the longitudinal immune changes of the TIME to assess the immune distance between the different timepoints of a same patient. A) Heatmap representation of the immune phenotype: hot and cold. Cold tumours, with overall low immune gene expression, are represented on the left of the graphic, whereas hot tumours, overexpressing most of the immune genes, are on the right. Each gene is represented in a row, and each patient in a column. Coloured labels above the graphic represent the different timepoints of a single patient. B) Alluvial flow diagram visualizing the evolution of the patients for their immune phenotype and immune gene expression clustering, defining the immune distance. C-D) Sankey flow diagrams of paediatric tumours indicated the tumour infiltrating-leukocyte populations reported on the Y-axis from highest (top) to lowest (bottom) cell density. The line width is scaled to cell density across timepoints *<u>Abbreviations</u>*: TIME: Tumour immune microenvironment, OtherST: others solid tumours, ES: Ewing sarcoma, STS: Soft-tissue sarcoma, RMS: Rhabdomyosarcoma, CD4+ Tem: CD4+ T effector memory, CD8+ Tem: CD8+ T effector memory, CMP: common myeloid progenitor, DC: dendritic cell, NK cells: natural killer cell, NKT: Natural Killer T cells, Tgd cells: Gamma delta T cell, Th1 cells: T helper 2 cells, Th2 cells: T helper 2 cells, Tregs: Regulatory T cell, D: diagnosis, R: relapse.

For immune gene expression clustering, the tumour samples clustered into five distinct centroids. Sixteen patients had all their timepoints grouped within the same centroid, while 11 patients had their timepoints spread across different centroids (Table 1).

Our immune distance evaluation, which incorporated the two metrics (immune phenotype and immune gene expression clustering), identified 13 patients as ‘distant’ and 14 as ‘close’ (Table 1, Supplementary Fig. S1). The immune changes or immune stability for patients within the two groups are summarized in an alluvial plot (Figure 1B). These findings support the hypothesis that immune gene expression undergoes temporal remodelling in a subset of paediatric tumours.

In addition to gene expression changes, immune cell infiltration also demonstrated temporal variability, further supporting the concept of TIME plasticity. This evolution was not linear, indicating dynamic modulation of the immune landscape. For example, two patients with immune evolution classified as distant showed significant modification of their immune cell infiltration during disease evolution. For patient 25, immune infiltration was principally composed of CD4+ T cells, natural killer (NK) cells, and Th1 cells in the first relapse, while the third relapse was mainly composed of M2 macrophages, natural killer cells and M1 macrophages, with broad changes observed across multiple immune cell types (Figure 1C). Similarly, patient 10 exhibited substantial shifts in immune cell composition over time. T cells and natural killer cells, initially present at high levels at the diagnostic, became one of the least abundant infiltrating populations at later timepoints. Conversely, the myeloid content became predominant with M2 and M1 macrophages highly represented in the fifth relapse (Figure 1D). When considering the entire cohort, no immune cell was found to be significantly modified. This precluded any definitive conclusion for overall trends in the evolution of tumour immune composition, which may instead reflect patient- or tumour-specific dynamics that were further explored in linear and time-dependent models.

### The immune landscape of paediatric tumours evolved toward a more tolerogenic environment over time

Generalized linear mixed models (glmm) for longitudinal differential gene expression analysis identified a total of 2250 genes significantly modulated over time, including 568 upregulated and 1682 downregulated genes (Figure 2A, Supplementary Table S2). Among these, 65 were immune-related genes, with 28 exhibiting increased and 37 decreased expressions over time. By assessing the gene function with UniProt of these immune genes, we revealed an evolution toward reduced anti-tumour activity and increased immunotolerogenic signalling. Upregulated genes were predominantly associated with tolerogenic cells, such as T regulatory cells and M2 macrophages: *TGFB3* (β = 0.27, q-value = 0.027; Figure 2B) (13), *CEBPD* (β = 0.27, q-value = 0.013; Figure 2C) (14), respectively. Other genes are inhibitors of pro-inflammatory immune cells like B cells or T cells: *TRIM55* (β = 0.15, q-value < 0.001; Figure 2D) (15), *NR4A1* (β = 0.20, q-value = 0.006; Figure 2E) (16), *CD276* (β = 0.14, q-value = 0.031; Figure 2F) (17). Conversely, downregulated genes were primarily involved in promoting anti-tumour immune responses, particularly T cell activation and function. These included *TMIGD2* (β = –0.20, q-value < 0.001; Figure 2G) (18), IL33 (β = – 0.28, q-value = 0.003; Figure 2H) (19), *DNTT* (β = –0.49, q-value < 0.001; Figure 2I) (20), and *IFNB1* (β = –0.13, q-value < 0.001; Figure 2J) (21).

**Figure 2.**
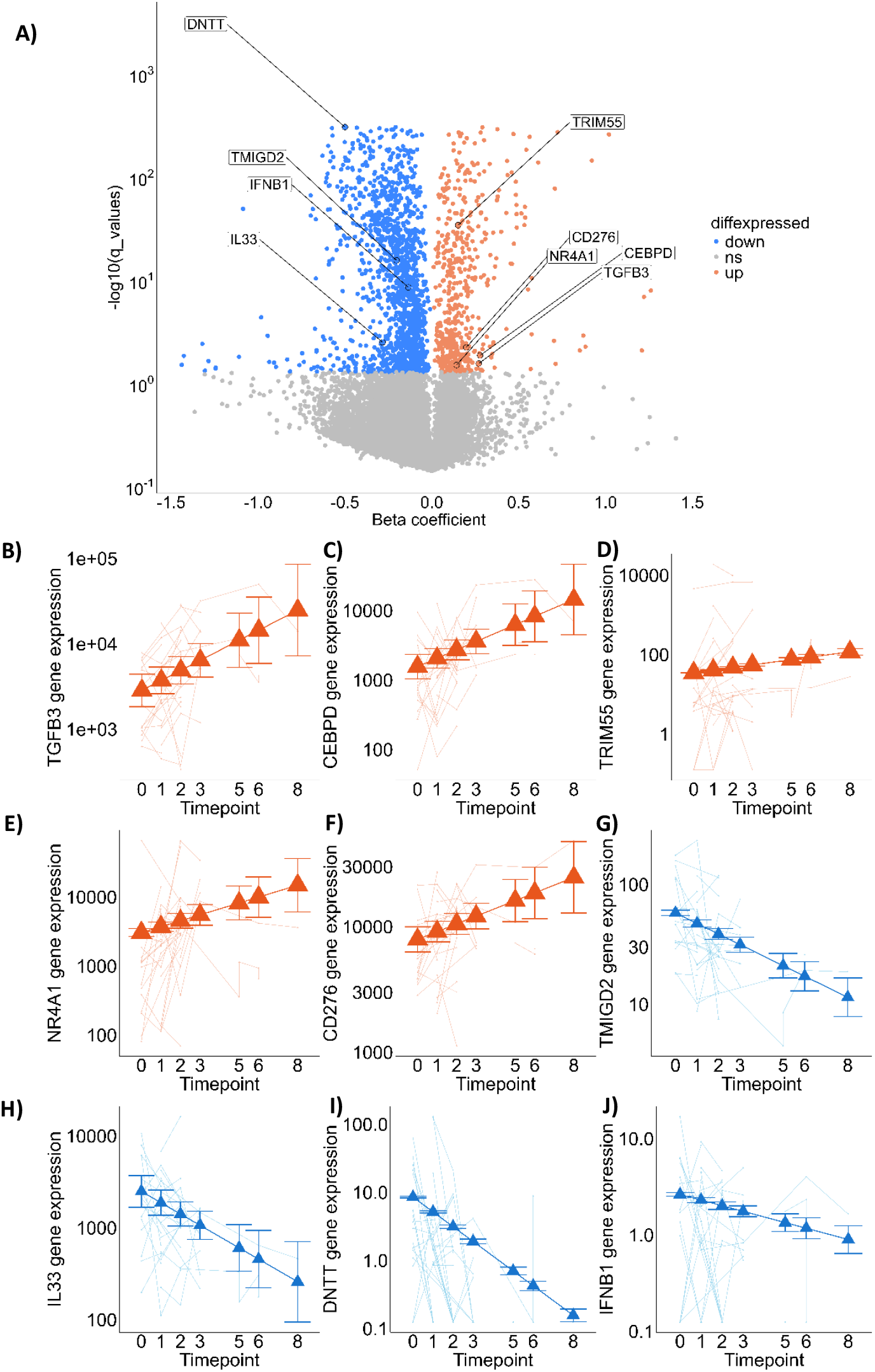
Representation of the immune gene’s evolution over time. A) Volcano plots of differentially expressed genes (DEGs) using glmmSeq comparing the evolution of the gene expression over time. Genes in blue are down-, and in red are up-regulated over time. The genes included in the panel of immune-related genes were annotated based on their role known in the bibliography. B-I) Scatter plots for selected DEGs with coloured points displaying the regression line of the fitted mixed-effects model. Error bars indicate 95% CIs (fixed effects). Orange and blue lines illustrate the evolution of the gene expression for each individual patient *<u>Abbreviations</u>*: CI: Confidence interval, q_values: adjusted p-value, ns: non-significant.

For longitudinal analysis, we also considered the potential impact of tissue origin—whether samples were collected from the primary tumour or metastatic sites. To address this, we performed a sensitivity analysis accounting for biopsy site, when available (15 patients in total). This analysis confirmed that, in our dataset, the biopsy site did not influence the observed changes in gene expression (Supplementary Fig. S2).

Collectively, these results suggest a progressive decline in anti-tumour immune activity and a concomitant increase in genes associated with immune tolerance, supporting the hypothesis that the TIME in paediatric tumours evolves toward a more immunosuppressive state over time.

### Tolerogenic pathways are enriched over time

Gene set enrichment analysis (GSEA) identified a total of 36 significantly enriched pathways from the HALLMARK database and 410 enriched and 16 depleted pathways from the Gene Ontology (GO) Biological Processes database (Table 1, Supplementary Table S3). Among the enriched HALLMARK pathways, four were directly related to immune regulation and tumour immune evasion: IL2/STAT5 Signalling (normalized enrichment score [NES] = 1.70, false discovery rate (FDR) < 0.001), IL6/JAK/STAT3 Signalling (NES = 2.17, FDR < 0.001), TGF-β Signalling (NES = 1.65, FDR = 0.003), and MYC Targets V2 (NES = 2.22, FDR < 0.001) (Figure 3A). From the GO database, two enriched immune-related pathways were identified: IL-6 production (NES = 1.57, FDR = 0.005) and macrophage differentiation (NES = 1.79, FDR = 0.007). Additionally, one immune-related pathway was significantly depleted: T cell receptor complex (NES = –1.79, FDR < 0.001) (Figure 3B).

**Figure 3.**
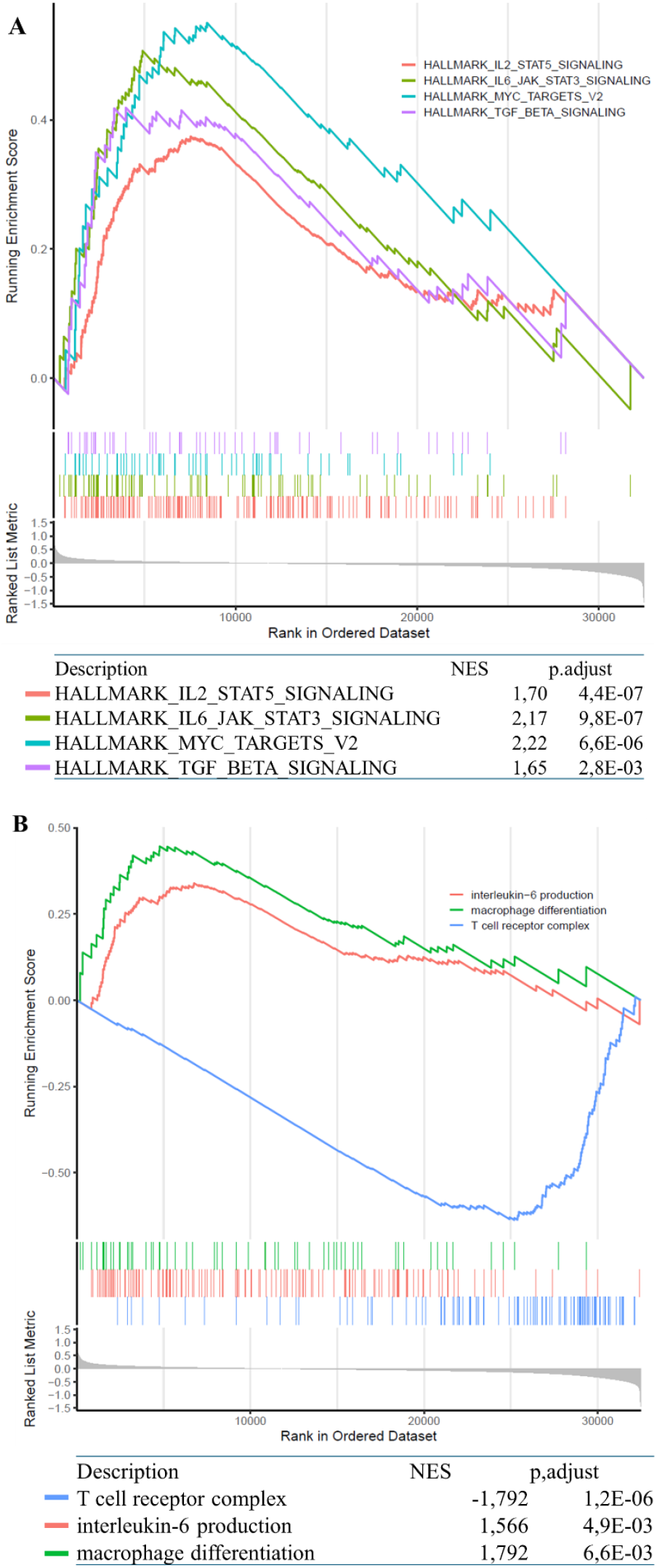
Gene set enrichment analysis identified a longitudinal enrichment of pro-tolerogenic pathways. A) Preranked gene set enrichment analysis using glmmSeq results in the hallmark database. B) Preranked gene set enrichment analysis using glmmSeq results in the GO database *<u>Abbreviations</u>*: p-adjust: adjusted p value, GO: gene ontology gene set library, NES: normalized enrichment score

Core enrichment analysis of top-expressed genes of these enriched pathways revealed several key immunoregulatory genes, including *CTLA4, IL10RA, TGFB1, CCL20, IL17RA, CSF1, CSF1R, CEBPB*, and *TRIB1*. These genes are known to contribute to immune suppression, T cell exhaustion, and the recruitment or polarization of immunosuppressive cell types such as regulatory T cells and M2 macrophages. Collectively, these findings suggest a progressive increase in TIME tolerogenicity and a concomitant reduction in the immune system’s capacity to eliminate tumour cells.

### Anti-tumour immune genes are correlated with oncogenic pathways

To explore the relationship between immune gene expression and oncogenic signalling, we performed weighted gene co-expression network analysis (WGCNA) using the 1,242 immune-related genes. This analysis identified a module—referred to as the green module—that was significantly correlated with two oncogenic HALLMARK pathways: MYC Targets V2 (correlation = 0.66, p-value < 0.001) and Hypoxia (correlation = 0.55, p-value < 0.001) (Figures 4A and 4B). The green module included immune genes such as tolerogenic immune genes, including *CCL20* (module membership = 0.74), *IL6R* (module membership = 0.83), and *DEF6* (module membership = 0.67), as well as the immune receptor genes *BTNL8* (module membership = 0.84) and *MST1R* (module membership = 0.93).

**Figure 4.**
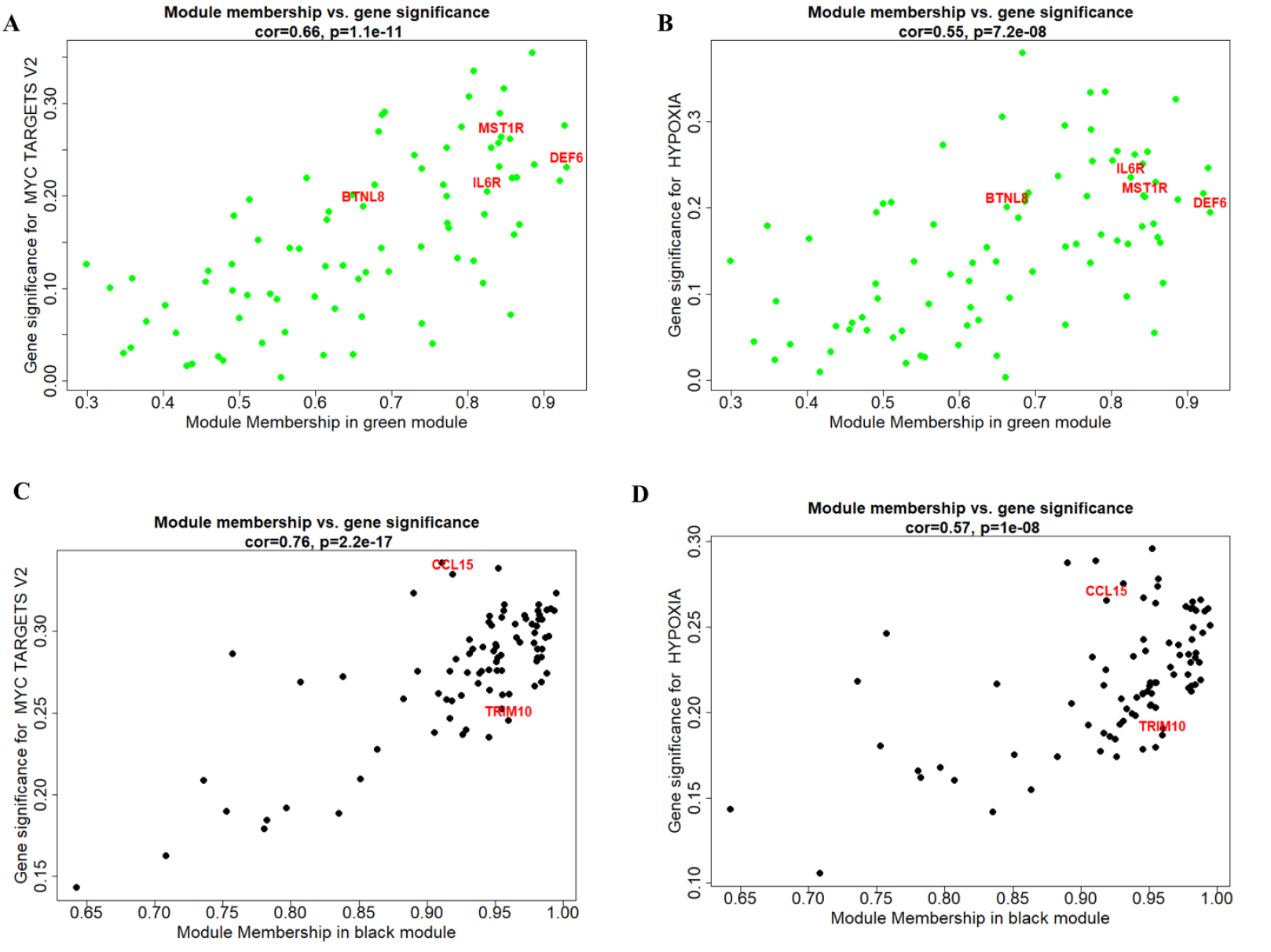
Network analysis identified immune genes correlating with oncogenic pathways. (A) Module membership in green represents genes highly correlated with MYC TARGETS V2 and (B) HYPOXIA. (C) Module membership in black displays genes with the highest coefficient of variation that correlate with MYC TARGETS V2 and (D) HYPOXIA. Highlighted genes were selected based on their association with immune-related functions. *<u>Abbreviations</u>*: cor: correlation, p: p-value.

A complementary WGCNA analysis was performed using the 4,000 most variable genes based on their coefficient of variation. This identified a black module that was also significantly correlated with MYC Targets V2 (correlation = 0.76, p-value < 0.001) and Hypoxia (correlation = 0.57, p-value < 0.001) (Figures 4C and 4D). The black module was predominantly composed of oncogenic genes but also included two immune-related genes: *CCL15* (module membership = 0.91) and *TRIM10* (module membership = 0.96). These findings indicate a positive correlation between immune gene expression and oncogenic pathway activity, suggesting that immune components may be co-regulated with tumour-promoting transcriptional programs during disease progression.

### Changes in immune genes and pathways are influenced by the time interval between timepoints

We hypothesized that the mechanisms driving immune changes within the TIME are influenced by the timing of relapse. Specifically, we propose that early relapses are predominantly shaped by the effects of prior treatments (extrinsic factors), whereas late relapses are more likely driven by intrinsic factors, such as tumour clonal evolution. Given that most chemotherapy regimens for paediatric cancers last 6 months or more, we defined early relapse as occurring within 180 days of the preceding timepoint, and late relapse as occurring beyond this threshold. To investigate whether specific immune genes and pathways were associated with the timing of tumour recurrence, we performed a recurrence analysis using a filtered set of differentially expressed genes (DEGs) and enriched immune pathways. Prior to analysis, highly correlated genes and pathways were removed to reduce redundancy. Of the 65 immune DEGs and 10 altered immune pathways initially identified, 49 genes and 6 pathways were retained for modelling. Using a recurrent event model, three genes were found to be significantly associated with shorter intervals between timepoints, suggesting a correlation with early relapses: *DNTT* (hazard ratio [HR] = 1.18, p-adjusted < 0.024), *CLCA1* (HR = 1.07, p-adjusted < 0.001), and *REG3G* (HR = 1.95, p-adjusted = 0.004). Conversely, *TGFB3* (HR = 0.92, p-adjusted < 0.001), *NR4A1* (HR = 0.99, p-adjusted = 0.034), and *CCL15* (HR = 0.98, p-adjusted = 0.034) were associated with longer intervals between timepoints, suggesting a reliance on intrinsic factors, such as tumour clonal evolution, rather than treatment-related influence. This observation aligns with the previously demonstrated correlation between *CCL15* expression and oncogenic signalling pathways. No immune-related pathways reached statistical significance (Figure 5, Supplementary Table S4 and S5).

**Figure 5.**
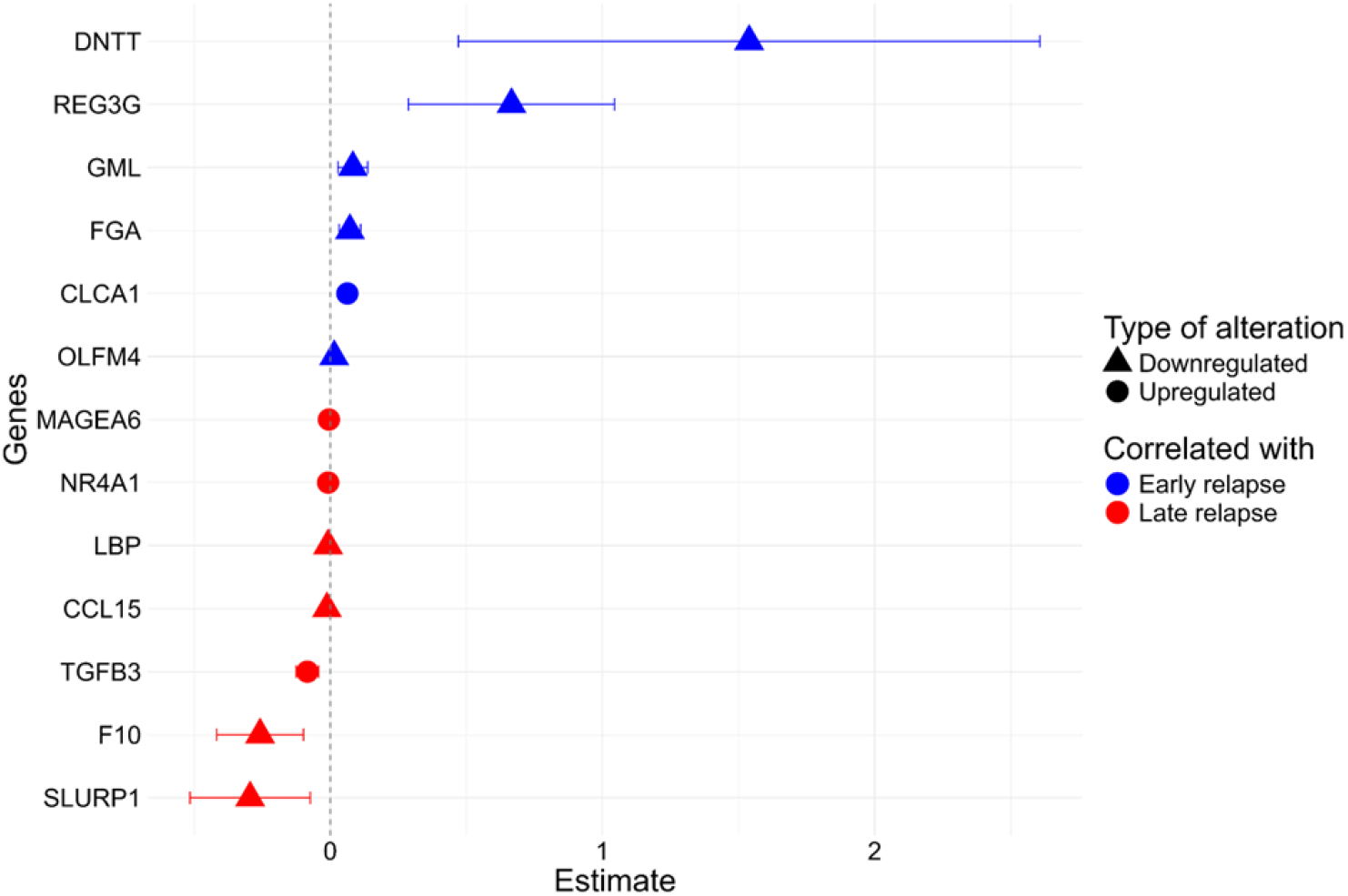
Time dependent immune genes in early (≤ 180 days) versus late relapses (> 180 days) Forest plot of significant correlation between differentially expressed genes and time between relapses was considered at p-adjusted <0.05. Blue indicates genes associated with early relapse, while red corresponds to those linked to late relapse. Circle-shaped markers represent upregulated genes, and triangle-shaped markers indicate downregulated genes.

In parallel, WGCNA was used to identify gene modules associated with the time between tumour sampling events. The lightgreen module demonstrated a significant correlation with time interval (correlation = 0.79, p-value < 0.001). Within this module, *IGHV3-64* was the only immune-related gene identified (module membership = 0.85), suggesting a positive correlation with the tumour clonal evolution.

## Discussion

Clustering based on immune gene expression revealed that paediatric solid tumours could transition between immune phenotypes—shifting from hot to cold or vice versa—across successive timepoints. This phenotypic plasticity was mirrored by changes in immune gene expression assessed by our immune distance metrics. Furthermore, immune infiltration levels were not stable over time, reinforcing the concept of TIME plasticity in paediatric solid tumours. The modifications of TIME indicate a shift toward increased immune tolerogenicity. This is evidenced by a downregulation of immune-related genes and an upregulation of pro-tumorigenic genes. Pathway analysis revealed an enrichment of tolerogenic signalling pathways, prominently involving *CSF1*/*CSF1R, MYC, TGFB1* and *IL10RA*—factors known to promote immune tolerance and support anti-inflammatory cell populations. The observed association between immune gene alterations and oncogenic pathway activation, along with the distinct timing of immune changes in early versus late relapses, suggests a dynamic interplay between tumour clonal evolution and immune remodelling.

These findings are consistent with previous observations in adult cancers, where TIME has been observed to evolve during disease progression and in response to therapy (10,22). In adult oncology, the development of tools such as CASCADE (Cancer Aggressiveness via Single-Cell Analysis During Evolution) has enabled the monitoring and prediction of TIME evolution. Using single-cell RNA sequencing, a scoring system was developed to classify patients’ TIME into four distinct categories. Following treatment, tumours could either retain their original classification or undergo significant changes. For instance, some patients initially exhibited a CD4^+^T cell–rich environment that later shifted toward increased infiltration by M2 macrophages and NK cells. The CASCADE framework demonstrated that the TIME can remain stable or be extensively remodelled during treatment, and importantly, that these transitions can be anticipated using CASCADE (23). Such predictive frameworks are instrumental in understanding how therapeutic interventions can alter the immune landscape and in guiding treatment decisions to improve clinical outcomes.

Our transcriptomic analysis revealed a progressive shift toward a tolerogenic TIME in paediatric cancers. This was characterised by the upregulation of key immunosuppressive genes and pathways, and the downregulation of genes associated with anti-tumour immunity. Among these, TGFB3 emerged as a central regulator, impairing T cell cytotoxicity and promoting exhaustion. In the TIME, activation of *TGFB* in effector T cells suppresses their cytotoxic activity and contributes to immune evasion (13). The enrichment of the TGFB signalling pathway, including *TGFB1* and *CTLA-4*, further supports this immunosuppressive shift. This pathway is well-documented for its role in dampening cytotoxic gene expression and facilitating tumour immune escape (24). Additional immunosuppressive mediators were also upregulated. *NR4A1* promotes T cell dysfunction (16). *CEBPD* contributes to a tolerogenic environment by reducing macrophage inflammatory responses and inducing IL6 expression—which is linked to immunosuppressive signalling (14,15). Which in turn activates the IL6/JAK/STAT3 pathway—a known driver of ICB resistance (25). Core enrichment genes such as *TGFB1, IL17RA*, and *TRIB1* were implicated in the upregulation of inhibitory surface molecules. TRIB1 is recognised as an anti-inflammatory regulator that maintains T cell exhaustion. Its deletion has been reported to restore effector gene expression, promote CD8+ T cell expansion, and enhance ICB efficacy (26–28). The enrichment of the IL2/STAT5 pathway further supports the immunosuppressive trajectory of the TIME. Persistent IL-2 signalling activates STAT5 in T cells, leading to increased expression of inhibitory receptors and reduced production of cytokines and effector molecules, ultimately rendering T cells dysfunctional (29). Core genes, such as *CTLA-4* and *IL10RA*—known to suppress T cell activation—were also enriched, reinforcing the immunosuppressive phenotype (30). Changes in the myeloid compartment were also observed. The enrichment of *CSF1* and *CSF1R* suggests increased polarisation toward M2 macrophages, which are associated with tumour progression and immune suppression (31–33). These findings advocate that the TIME becomes progressively more immunotolerant through the activation of tolerogenic pathways and the accumulation of anti-inflammatory myeloid cell populations. The existence of immune escape mechanisms that differ throughout disease evolution advocates for therapeutic strategies tailored to the immune evolution stages to foster the efficacy of immunotherapy for paediatric tumours.

Finally, the enrichment of oncogenic pathways, particularly MYC signalling, supports the notion that tumour-intrinsic factors contribute to TIME remodelling. MYC is a well-established oncogene that not only drives tumorigenesis but also modulates the immune microenvironment. It can drive immunotolerance and reshape the myeloid content toward tolerogenic cells (17,34). Correlation analysis using WGCNA revealed a strong association between *MYC* and immune gene expression. Coordinated regulation of oncogenic and immunosuppressive programs may collectively promote TIME tolerogenicity, providing insights for targeting strategies to reinvigorate anti-tumour immune activity.

Solving the longitudinal evolution of the tumour immune microenvironment in paediatric cancers requires the development of adapted analytical frameworks that are sorely lacking. To address the complexity of repeated measures and inter-patient variability, we employed a glmm approach using the glmmSeq package. This method is particularly well-suited for longitudinal transcriptomic data, as it accommodates more than two timepoints, handles missing data, and allows for the inclusion of covariates. Given the inherent heterogeneity of longitudinal cohorts—such as unbalanced sex and age distributions, and incomplete sampling across timepoints—this modelling strategy enabled us to extract meaningful biological signals while accounting for confounding variables. Sex and age are known to influence immune system function (35,36) and were therefore included as covariates in our model to ensure biological relevance. However, one limitation of our study was the inability to incorporate treatment regimens into the analysis, as treatment information was not uniformly available across patients. In our case, the biopsy site did not demonstrate a significant impact on the DEG. However, due to the high proportion of missing data and the fact that most biopsies were obtained from primary tumour sites, the biopsy location could still influence the results. Therefore, it remains important to consider the biopsy site as a potential confounding factor. Our findings were validated exclusively using the SIGNATURE dataset, due to the rarity of longitudinal tumour sampling and the current lack of publicly available longitudinal databases. Additionally, we did not perform cross-validation with orthogonal experimental approaches, which remains a limitation of this study. This study establishes a foundation for future research into the evolution of the tumour immune microenvironment through integrative multi-omics approaches. Integrating additional layers such as the methylome or proteome could provide a more comprehensive view of TIME dynamics and their concordance with transcriptomic changes. Spatial omics technologies, in particular, hold promise for elucidating the spatial organisation and evolution of tumour-infiltrating lymphocytes (TILs). These future directions will be essential for refining our understanding of TIME plasticity and for developing more precise, temporally informed immunotherapeutic strategies in paediatric oncology.

To conclude, our study represents the first longitudinal analysis of the tumour immune environment in paediatric solid tumours. Our findings revealed the dynamics of the TIME, highlighting a progressive shift towards a more tolerogenic immune landscape. These results underscore the critical importance of longitudinal sampling and comprehensive molecular profiling to inform the rational integration of ICB into therapeutic strategies for paediatric cancers.

## Materials and Methods

### Ethics statements

All participants provided written informed consent prior to their inclusion in the study. The Research Ethics Board at Centre Hospitalier Universitaire de Québec approved this study.

### Population

This study was conducted using the sequencing dataset from the SIGNATURE project, a prospective oncogenomic initiative of the province of Quebec aiming for molecular characterization of paediatric cancers (37). The database includes all types of paediatric malignancies, encompassing extracranial solid tumours, brain tumours, and hematologic cancers. For our longitudinal analysis, patients <21 years of age diagnosed with solid tumours—including both CNS and extracranial tumours—were considered. At the time of data cutoff, 434 patients with extracranial or CNS solid tumours were available. To ensure temporal resolution, only patients with at least two tumour samples collected at distinct timepoints were included in the dataset before the data cutoff on July 4^th^, 2024. The final dataset comprised 70 tumour samples from 27 paediatric patients.

### RNA sequencing workflow

Total RNA was extracted from the patient’s solid tumour cells using mini AllPrep DNA/RNA kits (80204) from Qiagen (QIAGEN RRID:SCR_008539). TruSeq Stranded Total RNA libraries were prepared using the Ribo-Zero Gold Kit (RS-122-2303) according to Illumina’s protocol (Illumina RRID:SCR_010233). The resulting libraries (stranded and ribosomal RNA-depleted) were sequenced (∼150M reads, principally paired-end 2×100bp, few 2×75bp for CNS tumour and some 2×150bp tumour) on either HiSeq 2500, HiSeq 4000 or NovaSeq 6000 systems at the Integrated Center for Clinical Pediatric Genomics at SJUHC, as previously described. (37).

Raw paired-end reads were trimmed using fastp (fastp, v0.22.0; fastp RRID:SCR_016962) (38). Pseudoaligned to the reference human transcriptome (Ensembl GRCh38 v100) using kallisto (kallisto, v0.48.0; kallisto RRID:SCR_016582) (39).

### Evaluation of the tumour immune plasticity

The concept of “immune distance” integrates two metrics evaluating immune changes: the immune phenotype and immune gene clustering. Both metrics were developed on a panel of 1,242 immune-related genes previously associated with tumour immune infiltration and ICB sensitivity (40–46). Patients switching either the immune phenotype or the immune cluster were considered “distant”, indicating a dynamic evolution of the TIME. In contrast, the “close” group includes patients whose tumours remain stable for these two metrics during disease evolution.

The immune phenotype used an in-house tumour immune classification to characterize the TIME of paediatric cancers within two immune phenotypes: “hot” and “cold”. Using an unsupervised classification with k-means (stats package, v 4.3.2; stats RRID:SCR_025968) trained on 340 tumour samples, tumours were classified based on the expression of immune-related genes. Samples exhibiting high expression of immune genes were classified as hot, indicative of an immune-infiltrated microenvironment, whereas those with low expression were categorized as cold. To identify immune subgroups, we performed hierarchical clustering (Ward.D2 method, stats package, v 4.3.2; stats RRID:SCR_025968) on the immune gene expression profiles of all 70 tumour samples collected across 27 patients with multiple recurrences. Importantly, this clustering incorporated all timepoints simultaneously, treating each sample as an independent observation regardless of its temporal position. This approach allowed us to assess how individual timepoints from each patient were distributed across clusters, thereby capturing temporal heterogeneity in immune profiles. The optimal number of clusters (centroids) was determined using Lloyd’s algorithm (stats, v 4.3.2; stats RRID:SCR_025968).

The immune distance was represented on an alluvial plot visualizing the evolution of the two metrics over time for each patient (ggalluvial, v 0.12.5; ggalluvial RRID:SCR_021253).

### Evolution of immune infiltrate

To investigate the temporal dynamics of immune cell infiltration, we estimated the abundance of immune cell populations using the xCell (v 1.1.0; xCell RRID:SCR_026446) algorithm (47), which infers cell type enrichment scores from transcriptomic data. Transcripts per million (TPM) values were used as input to xCell to quantify the relative enrichment of 20 immune cell types, including B cells, CD4+ T cells, CD4+ memory T cells, CD4+ effector memory T cells, CD8+ T cells, CD8+ effector memory T cells, common myeloid progenitors, dendritic cells, M1 and M2 macrophages, monocytes, NK cells, natural killer T (NKT) cells, γδ T cells, Th1 cells, Th2 cells, and Tregs. The xCell score for each cell type represents the average of its associated gene signature scores. Temporal changes in immune cell composition were visualized on alluvial plots (alluvial package in R, v 0.1-2, R package no RRID, Github: https://github.com/mbojan/alluvial). To identify immune cell types showing significant variation over time, we tested the xCell cell type scores using the Satterthwaite approximation (lmerTest, v 3.1-3, R package: lmerTest RRID:SCR_015656). Satterthwaite approximation is a method for estimating degrees of freedom in mixed models that improves the accuracy of statistical inference in this context. Multiple testing correction was performed using the Benjamini-Hochberg (BH) procedure, and significance was defined as a FDR < 0.05.

### Longitudinal analyses of gene expression

The objective of this analysis was to identify immune-related genes that are differentially expressed over time and represent the evolution of the TIME. To this end, a longitudinal RNA-seq analysis was performed using a glmm framework, implemented via the glmmSeq (glmmSeq, v 0.5.5; R packages no RRID, Github: https://cran.r-project.org/web/packages/glmmSeq/index.html) R package (48). This approach allows for the modelling of gene expression evolution, using raw counts, across multiple timepoints while accounting for inter-patient variability and missing data— limitations that are not addressed by conventional differential expression tools. For each gene, a negative binomial glmm was fitted using the glmer() function from the lme4 package (lme4, v1.1-25, lme4 RRID:SCR_015654), in combination with the negative.binomial() function from the MASS package (MASS, v7.3-53; Modern Applied Statistics with S RRID:SCR_019125). Raw count data were pre-processed by filtering out lowly expressed genes using the filterByExpr() function with default parameters. To ensure model stability, persistent zero counts were replaced with a small constant value (zeroCount = 0.125). Model fitting was performed using maximum likelihood estimation via Laplace approximation and bound optimization by quadratic approximation. Timepoint was included as a fixed effect, while patient ID, sex, and age were treated as a random effect to account for repeated measures. Sensitivity analyses also tested the effect of tumour sites (primary versus metastasis). Multiple testing correction was applied using the q-value method, and genes with an FDR ≤ 0.05 were considered significantly differentially expressed.

### Longitudinal gene set enrichment analyses

To identify immune-related pathways involved in the longitudinal evolution of the TIME, GSEA was performed using the results from the glmmSeq differential expression analysis. Genes were ranked based on the β-coefficient values derived from the glmm models, reflecting the direction and magnitude of expression changes over time. Ranked gene lists were used as input for GSEA, implemented via the clusterProfiler R package (clusterProfiler, v 4.10.1; clusterProfiler RRID:SCR_016884). Enrichment analysis was conducted using two reference databases: the GO and the HALLMARK gene sets. For the GO we selected the biological process (BP), cellular component (CC), or molecular function (MF). Pathways with FDR (BH) ≤ 0.05 were considered significantly enriched.

### Co-expression network analyses

To investigate the interplay between tumour cell oncogene activation and immune remodelling, we explored the strength of association between immune gene expression and oncogenic pathways. We performed gene module association using WGCNA, as implemented in the WGCNA R package (WGCNA, v 1.72-5; Weighted Gene Co-expression Network Analysis (RRID:SCR_003302)). The analysis was first conducted on the 1,242 immune-related genes previously used for immune distance analysis. Gene modules were identified using the blockwiseModules() function, with an unsigned network constructed using a soft-thresholding power of 3, as determined by the pickSoftThreshold() function. Oncogenic pathway activity scores were treated as external clinical traits and correlated with the resulting gene modules. Modules significantly associated with pathway activity were further explored to identify the immune genes they contained and to characterize their functional relevance.

A complementary WGCNA analysis was performed using the 4,000 most variable genes, selected based on their coefficient of variation across all samples. The network was constructed using the same blockwiseModules() function, with an unsigned network and a soft-thresholding power of 27, also determined via pickSoftThreshold(). Modules significantly correlated with oncogenic pathways were similarly analysed to identify immune gene content and to explore their potential role in TIME evolution.

### Time-dependent immune changes in early and late relapses

To identify genes and pathways associated with early (≤180 days from previous timepoints) versus late relapses (>180 days from previous timepoints) (49), we first filtered the list of DEGs, and enriched pathways obtained from the glmmSeq and GSEA analyses to reduce redundancy and background noise. A Spearman correlation analysis was performed to identify highly correlated genes and pathways. For gene pairs with a correlation coefficient > 0.7 for raw counts, one gene from each pair was removed to reduce redundancy. If a gene was paired with multiple other genes, the genes with the highest number of high-correlation pairings were iteratively removed until no pairs exceeded the 0.7 threshold. Similarly, for pathways with a correlation coefficient > 0.7, gene set overlap was assessed. If pathways shared more than 10% of their constituent genes, one was excluded from further analysis. The remaining genes and pathways were then analysed using a recurrent event model to assess their association with the time interval between two events. This analysis was performed using the TimeReg R package (TimeReg, v 2.0.5; R package no RRID, CRAN: http://dx.doi.org/10.32614/CRAN.package.timereg) (50), which implements the Ghosh and Lin model for recurrent event data. The model evaluated the relationship between gene/pathway expression and the time interval between tumour sampling timepoints. Genes or pathways with an estimate greater than 0 were associated with early relapse, whereas those with an estimate less than 0 were linked to late relapse. Significance was reached with an FDR (BH) ≤ 0.05.

In parallel, we used WGCNA to assess whether modules of DEGs were associated with the time interval between events. The differentially expressed genes were used as input for network construction via the blockwiseModules() function. An unsigned network was generated using a soft-thresholding power of 12, selected using the pickSoftThreshold() function. Modules significantly associated with the time separating two timepoints were further analysed to identify immune-related genes and to characterize their potential role in TIME evolution.

## Supporting information

Supplementary Fig. S1: Clustering of patients to measure the immune distance

Supplementary Fig. S2: Sensitivity analysis to test the effect of tumour sites (primary versus metastasis) on the DEGs

Supplementary description

## Data availability statement

The transcriptomic data will be accessible through dbGaP (NCBI database of Genotypes and Phenotypes (dbGap) RRID:SCR_002709) repository and available upon request.

## Package availability

The R package glmmSeq is downloadable via CRAN, and the source code is also available, from https://github.com/myles-lewis/glmmSeq.

The package alluvial is downloadable via the source code is also available, from https://github.com/mbojan/alluvial

The package WGCNA is downloadable via the source code is also available, from https://cran.r-project.org/web/packages/WGCNA/index.html.

## Acknowledgments

I thank all the contributors to this article.

