## Supplementary figures and images for "Unveiling the temporal impact: Exploring dynamic changes in the paediatric solid tumour immune microenvironment through time"

### Supplementary Fig. S1: Clustering of patients to measure the immune distance

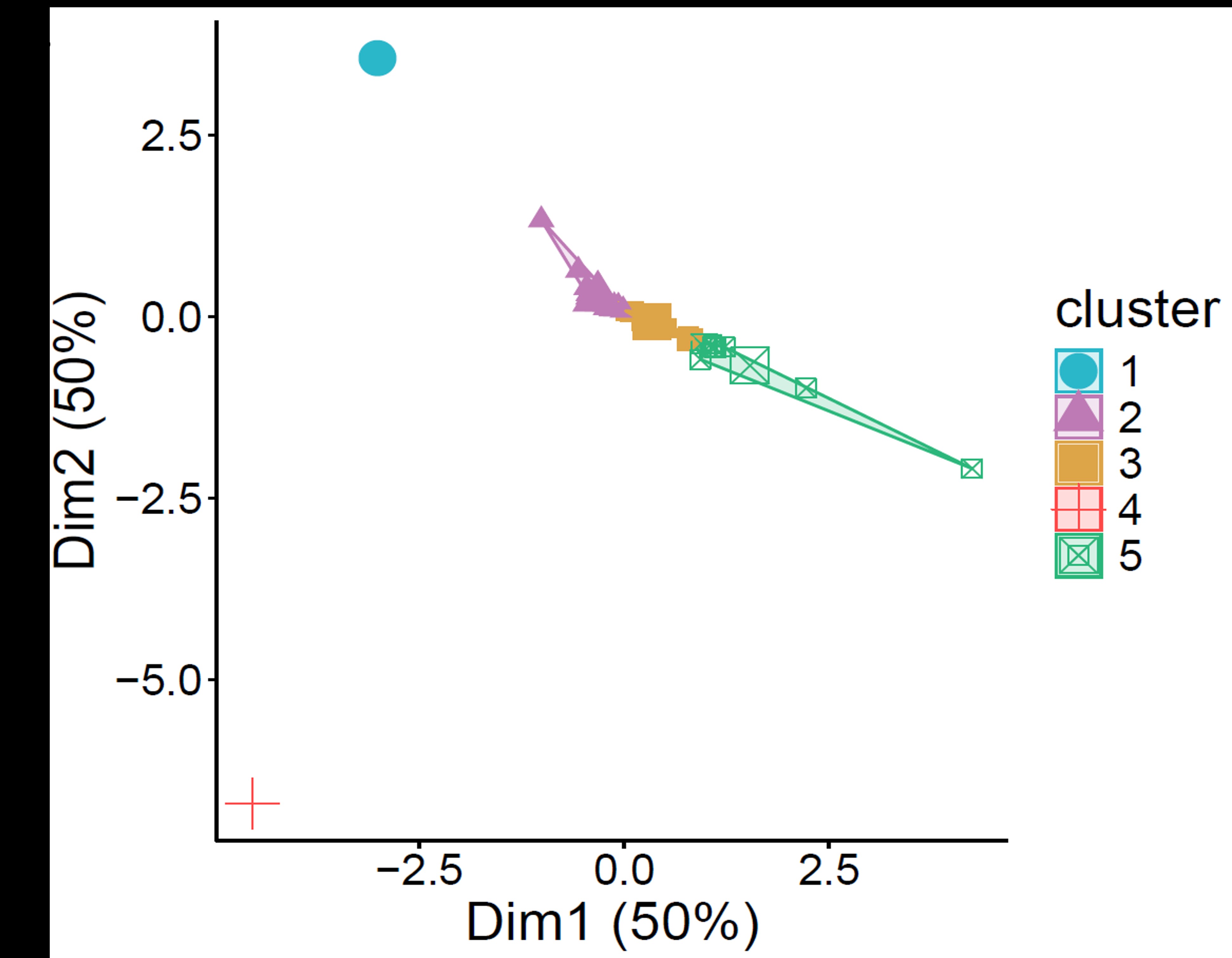

### Supplementary Fig. S2: Sensitivity analysis to test the effect of tumour sites (primary versus metastasis) on the DEGs

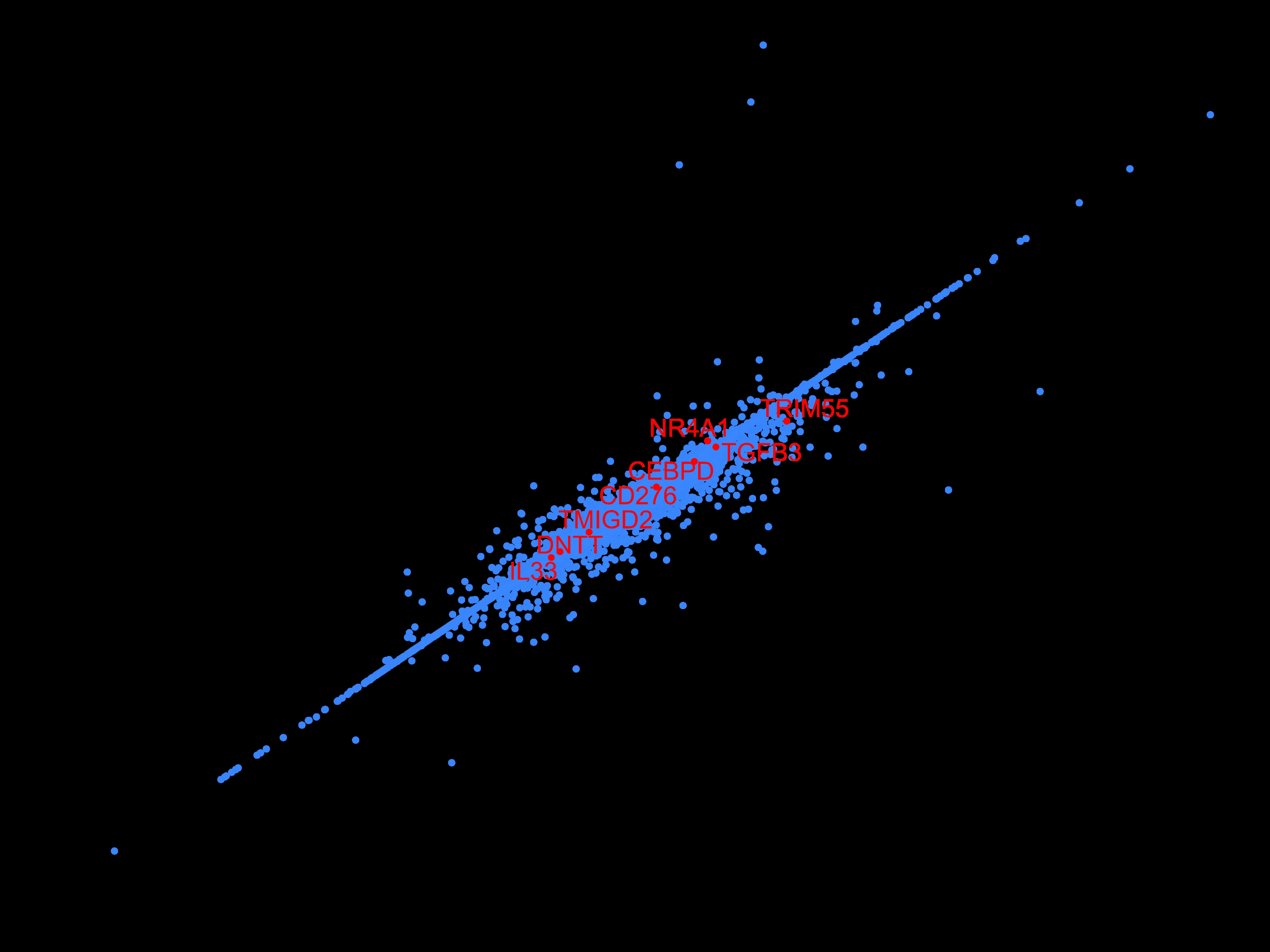
