## Supplementary description for "Unveiling the temporal impact: Exploring dynamic changes in the paediatric solid tumour immune microenvironment through time"

**SUPPLEMENTARY DATA:**

**Table S1. Summary of the patients’ characteristics, describing the demographics, age at the time of biopsy, tumour type, time between biopsies and site of the biopsy**

Abbreviations: *D*: diagnostic, *R*: relapse, *Patient ID*: Patient identification

**Table S2. Data frame of the longitudinal DEG analysis**

The table includes the results of all the genes analysed in the DEG analyses. Dispersion, mean expression, beta-coefficient, *p*-value, and q-value are displayed in the data frame to summarise the results of the analysis. Significant values are in bold.

Abbreviations: *AIC*: Akaike’s Information Criteria, *se_(Intercept)*: standard-error intercept, *meanExp*: mean expression, Chisq_*Time*: Wald Chi-square test on beta, *se_Time*: standard-error of the beta-coefficient

**Table S3. Results of the gene set enrichment analysis table composed of Hallmark, GO and KEGG**

Display the results of the gene set enrichment analysis for the three libraries: hallmark, gene ontology (GO) and Kyoto encyclopedia of genes and genomes (KEGG). Only significant data are included.

Abbreviations: *p.adjust*: adjusted *p* value, *GO*: gene ontology gene set library, *NES*: normalized enrichment score, *setSize*: gene set size, *ID*: identifications, *KEGG*: Kyoto encyclopedia of genes and genomes

**Table S4. Data frame of the TimeReg analysis for the DEG**

Result of the recurrent event analysis using TimeReg. Each row displays a gene that has been tested for correlation with early or late relapse. The estimates with positive values are associated with early relapses, and negative values with late relapses. The significance was achieved at 0.05 for the adjusted *p*-value. Significant values are in bold.

Abbreviations: *SE*: standard-error, *P_adjusted:* adjusted *p*-value, dU: derivative of the score function

**Table S5. Data frame of the TimeReg analysis for the pathways**

Result of the recurrent event analysis using TimeReg. Each row displays a pathway tested to be correlated with early or late relapse. The correlation with early relapse was deducted if the estimate result was positive, negative values were associated with late relapse. The significance was achieved at 0.05 for the adjusted *p*-value

Abbreviations: *SE*: standard-error, *P_adjusted:* adjusted *p*-value, dU: derivative of the score function

**SUPPLEMENTARY FIGURE LEGENDS**

**Supplementary Fig. S1: Clustering of patients to measure the immune distance**

**LEGEND**: Clustering of the immune distance using Ward.D2 linkage method, based on 1242 immune genes. The optimised number of clusters was determined using Lloyd’s algorithm. Each dot represents a timepoint for a patient

Abbreviations: *Dim*: dimensions

**Supplementary Fig. S2: Sensitivity analysis to test the effect of tumour sites (primary versus metastasis) on the DEGs**

**LEGEND**: Comparison of genes’ fold change when tumour site, age and sex were used as fixed effects (axis x) compared to age and sex adjustment only (axis y)

Abbreviations: *Site*: biopsy site, *FC*: Beta-coefficient

**SUPPLEMENTARY FIGURE:**

**Figure S1:**

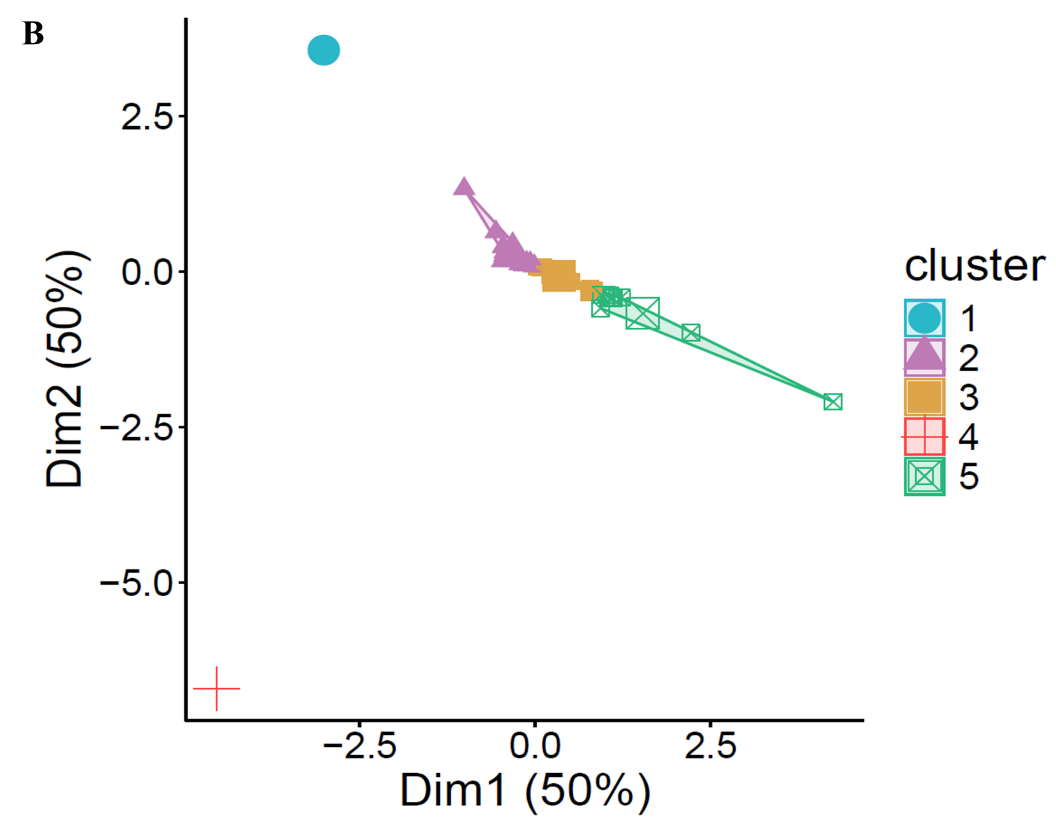

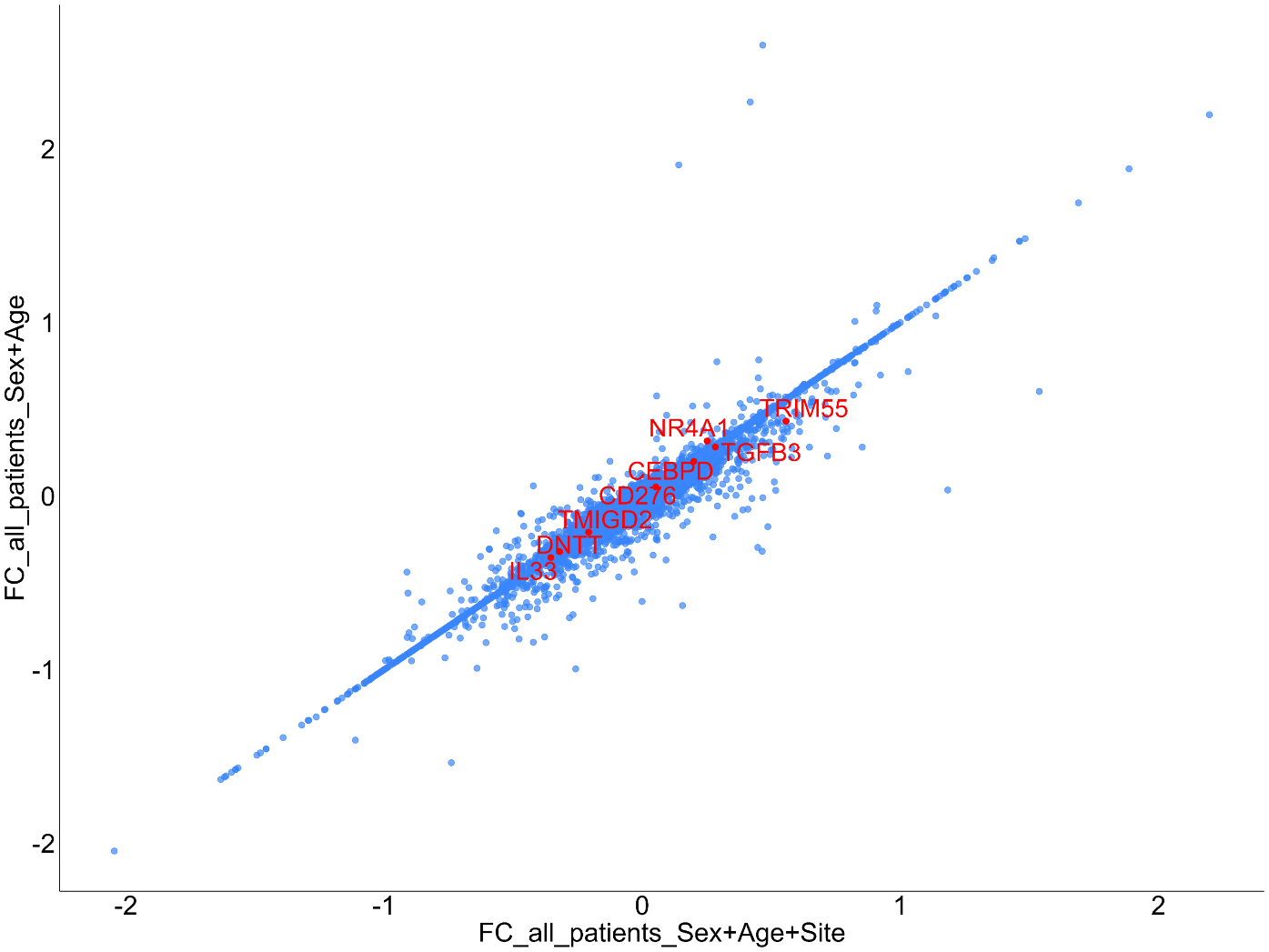
**Figure S2:**
